# In vivo characterization of neurophysiological diversity in the lateral supramammillary nucleus during hippocampal sharp-wave ripples of adult rats

**DOI:** 10.1101/712596

**Authors:** Ana F Vicente, Andrea Slézia, Antoine Ghestem, Christophe Bernard, Pascale P Quilichini

**Affiliations:** Aix Marseille Univ, Inserm, INS, Institut de Neurosciences des Systèmes, Marseille, France

## Abstract

The extent of the networks that control the genesis and modulation of hippocampal sharp wave ripples (SPW-Rs), which are involved in memory consolidation, remains incompletely understood. Here, we performed a detailed *in vivo* analysis of single cell firing in the lateral supramammillary nucleus (lSuM) during theta and slow oscillations, including SPW-Rs, in anesthetized rats. We classified neurons as SPW-R-active and SPW-R unchanged according to whether or not they increased their firing during SPW-Rs. We show that lSuM SPW-R-active neurons increase their firing prior SPW-Rs peak power and prior hippocampal pyramidal cell activation. Moreover, lSuM SPW-R-active neurons show increased firing activity during theta and slow oscillations as compared to unchanged-neurons. SPW-R-active neurons are more active during high peak power SPW-Rs, whereas SPW-R-unchanged neurons are more active during long SPW-Rs. These results suggest that a sub-population of lSuM neurons can interact with the hippocampus during SPW-Rs, raising the possibility that the lSuM may modulate memory consolidation.

## Introduction

Sharp-wave ripples (SPW-Rs) consist of a distinctive oscillation of the hippocampal local field potential (LFP) in the CA1 region that can occur during sleep and awake immobility (Buzsáki, 2015). This phenomenon consists of two components: a negative deflection called sharp wave and a fast oscillation (150-250Hz) called ripple. Several studies suggest that SPW-Rs play a key role in the transfer of information between the hippocampus (HPC) and the neocortex (Diekelmann & Born, 2010; Maingret et al., 2016), but the mechanisms underlying the genesis of SPW-Rs remain mostly unknown. They appear to be an emergent property of neuronal networks, which could involve different brain regions, as for gamma oscillations (Buzsáki & Schomburg, 2015; Sohal & Huguenard, 2005). Several studies have shown the involvement of the CA2/CA3 subfields of the HPC (Jones & McHugh, 2011). CA3 synchronous bursts depolarize CA1 pyramidal cells in stratum radiatum, resulting in a negative LFP deflection (sharp waves) (Buzsáki *et al.*, 1983). This synchronous activity triggers the firing of CA1 interneurons at a high frequency (Csicsvari *et al.*, 2000). The inhibition of interneurons together with CA3 activation result in fast field oscillations known as ripples (Ylinen *et al.*, 1995).

A change in firing frequency around SPW-Rs is generally considered as a marker of their involvement in SPW-R genesis and/or in memory consolidation process. For example, HPC neurons increase their firing during SPW-R events in sleep, reflecting neuronal patterns associated with previous experiences (Lee & Wilson, 2002). Oliva *et al.* (2016) demonstrated that CA2 neurons firing precedes the activation of CA3 and CA1 during SPW-Rs, suggesting a prominent role of CA2 region in SPW-R genesis. In line with a distributed network phenomenon, a number of studies have looked at the interaction between SPW-Rs and different subcortical areas (Girardeau *et al.*, 2017; Pennartz *et al.*, 2004). A typical example is the septum, whose neurons increase or decrease their discharge probability during SWP-Rs (Dragoi *et al.*, 1999; Unal *et al.*, 2018). A clear understanding of the mechanisms underlying the genesis of SPW-Rs may require the identification of all the structures that can control or modulate HPC SPW-Rs.

The goal of the present study is to determine whether the supramammillary nucleus (SuM) is part of the network that can generate or modulate SPW-Rs, using as a marker a change in firing activity of SuM cells around the time of occurrence of HPC SPW-Rs. The SuM is a hypothalamic region divided in medial and lateral parts (lSuM) according to their respective different connections to the HPC (Pan & McNaughton, 2004). The lSuM sends projections to the dentate gyrus throughout both dorsal and ventral HPC, to dorsal CA2/CA3 (Vertes, 1992), and is reciprocally connected with the septal complex (Borhegyi *et al.*, 1998; Leranth & Kiss, 1996). Electrophysiological studies of the SuM showed its role in theta oscillation (THE) due to its connections with the septum, which is considered a THE pacemaker (Pignatelli *et al.*, 2012). Kocsis & Vertes (1994) described SuM neurons firing rhythmically with THE recorded in dorsal HPC. Moreover, Kirk & McNaughton (1993) showed that pharmacological blockade of the SuM results in a decrease of frequency and amplitude of hippocampal THE. These results suggest the involvement of the SuM in THE modulation. However, no studies have described the activity of the SuM during SO and its possible role in SPW-Rs. The connectivity of the SuM with both HPC and septum (Borhegyi *et al.*, 1998; Kiss *et al.*, 2000) provides a morphological substrate for a potential involvement of the lSuM in the control/genesis of SPW-Rs.

To examine this possible involvement during SO, 5 Wistar rats were implanted with silicone probes into dorsal CA1 HPC and lSuM. This procedure allowed us to record simultaneously LFP and firing activity from the two regions during both THE and SO under anesthesia. We found a population of lSuM neurons increasing its firing around hippocampal SPW-Rs, which we called “SPW-R-active neurons”. Additionally, these neurons showed distinct firing properties comparing to those lSuM neurons showing no firing increases during SPW-Rs.

## Material and Methods

### Ethical approval

All experiments were performed in accordance with Aix-Marseille Université and Inserm Institutional Animal Care and Use Committee guidelines. The protocol was approved by the French Ministry of National Education, Superior Teaching, and Research, approval number (01451-02). All surgical procedures were performed under anesthesia. All experiments described here comply with the policies and regulations described in Grundy (2015)

### Animal surgery

5 Wistar Han IGS male rats (Charle Rivers) were used in this study. Animals were housed for at least 7 days upon arrival in groups of 2 in a temperature (22 ± 1 °C) and humidity (60~70%) controlled facility on a 12 h light-dark cycle (07.00 to 19.00 h) with food and water available *ad libitum*. Animals were anesthetized with urethane (1.5 g/kg, i.p.) and ketamine/xylazine (20 and 2 mg/kg, i.m.) with additional doses of ketamine/xylazine (2 and 0.2 mg/kg) when needed during the electrophysiological recordings. Animal breathing, hear rate, pulse distension, and arterial oxygen saturation were monitored with an oximeter (MouseOX; StarrLife Science) during the whole experiment. The animal was placed in a stereotaxic frame, its head was exposed. Two stainless-steel screws were implanted into the skull above the cerebellum, serving as reference and ground electrodes. Two craniotomies were performed to target the dorsal CA1 HPC (coordinates from bregma AP −3 mm, ML −2.4 mm, DV −2.4 mm, with 20° angle) and the lSuM (coordinates from bregma AP −4.44 mm, ML −2 mm, DV −8.6 mm, with 10° angle). HPC and lSuM activity were simultaneously recorded with 32-site silicon probes (Neuronexus Technologies). lSuM was recorded with an H32-Edge-10mm-20 μm-177 probe for all animals, and HPC was recorded with the following probes: an H32-Poly2-5mm-25μm-177 probe for animal A, a linear H32-6mm-50μm-177 probe for animal B, a linear H32-10mm-100μm-177 probe for animal C, and a two-shank linear H16x2-10mm-100μm-177 probe for animals E and F.

All silicon probes were mounted on individual stereotaxic manipulators and lowered manually for HPC and with a motorized descender (IVM, Scientifica) for lSuM. The final position of the probes was adjusted according to the presence of unit activity in cell body layers and the presence of ripples [100 200] Hz in CA1 stratum pyramidale.

### Electrophysiological recordings and initial analysis

Extracellular signal was amplified (1000X), bandpass filtered (1 Hz to 5kHz) and acquired continuously at 32kHz with a Digital Lynx (NeuraLynx) at 16-bit resolution. Raw data were preprocessed using NEUROSUITE (Hazan *et al.*, 2006; http://neurosuite.sourceforge.net/; RRID: SCR_008020). The signal was downsampled to 1250 Hz for the LFP analysis. Spike sorting was performed automatically, using KLUSTAKWIK (Harris *et al.*, 2000; http://klustakwik.sourceforge.net/; RRID: SCR_014480), followed by a manual adjustment of the clusters, with the help of auto-correlogram, cross-correlogram, and spike wave-form similarity matrix with KLUSTERS software (Hazan *et al.*, 2006; RRID: SCR_008020). Only units showing a clear refractory period and a well-defined cluster were included for posterior analysis. Theta and SO periods were identified for each recording. LFP THE or SO epochs were visually selected from the ratios of the whitened power in the THE band (3-6 Hz) and the power of the neighboring bands (1-3, and 7-14 Hz) of CA1 pyramidal layer, and from the ratio of the whitened power in the SO band (0.5-2 Hz) of CA1 pyramidal layer, respectively. This procedure was assisted by visual inspection of the raw traces (Quilichini *et al.*, 2010).

### Single unit identification

Units were identified as lSuM neurons by determining the histological reconstruction of the recording silicon probes tracks, the approximate location of their somata relative to the recording sites, and the known distances of the recorded sites. Hippocampal putative pyramidal neurons and interneurons were separated on the basis of their auto-correlograms, firing rates, and waveforms features, assisted by monosynaptic excitatory and inhibitory interactions between pairs of neurons simultaneously recorded reflected on the cross-correlograms (Barthó *et al.*, 2004; Csicsvari *et al.*, 1998).

### Histological analysis

At the end of the recording, the animals were injected with a lethal dose of pentobarbital Na (150 mg/kg, i.p.) and perfused intracardially with 4% paraformaldehyde solution in 0.12 M phosphate buffer (PB), pH 7.4. The brains were removed and post-fixed at 4°C overnight. They were then rinsed in PB and sliced into 60-μm-thick coronal sections by a Vibratome. The position of the electrodes was revealed by the presence of DiIC18(3) (Interchim), which was applied on the back of the electrodes before insertion and confirmed histologically on fluorescent Nissl-stained sections (Neuro-Trace 500/5225 Green Fluorescent Nissl Stain, Invitrogen). Only experiments with the appropriate position of the electrodes were used for analysis.

### SPW-Rs analysis

SPW-Rs were detected during SO periods with RippleLab (Navarrete *et al.*, 2016) independently on each recording. LFP corresponding to the hippocampal stratum pyramidale was digitally band-pass filtered 80-250 Hz, and the power (root-mean-square) of the filtered signal was calculated. The mean and SD of the power signal were calculated to determine the detection threshold. Oscillatory epochs with a power of 5 or more SD above the mean were detected. The beginning and the end of oscillatory epochs were marked at points where the power fell <0.5 SD. Short events (>15ms) were discarded and adjacent events (gap>15ms) were merged. All events were visually confirmed to avoid false positives. Only those events simultaneously detected with a sharp wave in the stratum radiatum were kept for further analysis, confirmed by the negative correlation between the whiten signal of each ripple event and its corresponding sharp wave, assisted by visual inspection. Inter-SPW-Rs periods were defined as SO periods by excluding SPW-Rs times.

### Correlation of unit firing by SPW-Rs

Spikes of each neuron detected in a 200 ms window centered on the peak of each ripple power were binned (5 ms time bin) to construct ripple cross-correlograms. To assess the significance of these cross-correlograms a nonparametric significance test based on jittering was used (Amarasingham *et al.*, 2012; Ferraris *et al.*, 2018; Fujisawa *et al.*, 2008). Surrogate test (n=1000 surrogates, 100ms window interval) was applied to set the 95% upper and lower confidence intervals. Then, the cross-correlograms were constructed for the surrogate datasets as a function of latency across the interval −100 +100 ms. Pointwise line at 99% acceptance level were constructed for the cross-correlograms from the maximum and minimum of each jitter surrogate cross-correlogram across the interval −100 +100 ms. Neurons crossing the upper or lower pointwise lines for at least 3 consecutive bins were considered as positively or negatively correlated by SPW-Rs, respectively.

Raster plots were constructed for each neuron by collecting the spikes detected in a 400 ms window centered on the peak of each ripple power. Spikes were binned (1 ms time bin) and cross-correlograms were constructed. The cross-correlograms were z-scored and smoothed using a Gaussian kernel (SD=5 ms) to construct the raster plots. The latency peak firing during SPW-R for each neuron was based on these z-scored smoothed cross-correlograms. For each type of neuron (pyramidal, interneurons and lSuM neurons), all z-scored smoothed cross-correlograms were pooled and mean and SD curves calculated.

Participation probability was calculated for each neuron defined as the number of SPW-Rs in which a neuron fires divided by the total number of SPW-Rs (Oliva *et al.*, 2016).

### Firing properties analysis

Firing properties were analyzed for each neuron to determine differences among the types of neurons for both THE and SO.

Bursting was analyzed by calculating a burst index for each neuron obtained from the autocorrelogram of each neuron (1 ms bin) by subtracting the mean frequency between 40-50 ms (baseline) from the maximum frequency between 0-10 ms. Positive values were normalized to the peak frequency and negative values were normalized to the baseline to obtain indexes ranging from −1 to 1. Indexes >=0.6 indicated bursting. Intra-burst frequency was defined as the maximum frequency between 0-10 ms and was calculated for those neurons showing a burst index>=0.6.

We also analyzed the firing properties for every SuM neuron during SPW-R and inter-SPW periods (Katona *et al.*, 2014; Unal *et al.*, 2018). SPW-R firing rate was calculated by summing all spikes during the detected SPW-Rs and dividing the result by the sum of durations of all SPW-Rs. To calculate firing rate for inter-SPW-R epochs, a set of 1000 surrogates time windows (surrogate SPW-Rs) was created for every SPW-R. Surrogate SPW-Rs were restricted to SO periods by excluding SPW-R periods. The firing rate for every surrogate SPW-R was calculated and its average was derived for every neuron. The same procedure was followed to calculate burst index and intra-burst frequency.

Autocorrelograms (1 ms bin size) were built for every neuron for THE and SO periods.

### Statistics

All results reported are based on estimation statistics which provide the effect size and the 95% confidence interval (CI) of the median difference between two groups. We directly introduced the raw data in https://www.estimationstats.com/ and downloaded the results and graphs. The median difference for two comparisons is shown with Cumming estimation plot. The raw data is plotted on the upper axes. For each group, summary measurements (mean ± standard deviation) are shown as gapped lines. Each median difference is plotted on the lower axes as a bootstrap sampling distribution. Five thousand bootstrap samples were taken; the confidence interval was bias corrected and accelerated. Median differences are depicted as dots; 95% confidence intervals are indicated by the ends of the vertical error bars. In order to measure the effect size, we used the difference between medians. We also provide P value(s) the likelihood(s) of observing the effect size(s), if the null hypothesis of zero difference is true, using the test mentioned in the text. If 95% CI includes 0, differences are considered as non-significant. P values are also provided. In the case 95% CI includes 0 and p-values is lower than 0.05, differences are also considered as non-significant. We used nonparametric testing in most cases: two-sided paired testing Wilcoxon’s signed-rank test for paired groups and two-sided Kruskal test for unpaired groups and provided the median value for each group.

To establish the THE and SO modulation of units, the phase of the hippocampal slow oscillations was determined from the filtered LFP in the (3-6 Hz) and 0.5-2 Hz, respectively. The instantaneous phase was computed as the angle of the Hilbert transform. Using linear interpolation, a value of phase was assigned to each spike. The modulation of lSuM spikes was determined by Rayleigh circular statistics. P values<0.05 were considered significant.

## Results

### Neuronal firing during SPW-Rs

We recorded simultaneously LFP and single unit activity from dorsal CA1 HPC and lSuM with silicon probes in anesthetized rats during theta (THE) and slow oscillations (SO) epochs. Hippocampal neurons were separated into putative pyramidal neurons and interneurons. SPW-Rs occurring during SO were detected. We classified neurons as a function of the way their firing activity was changed during SPW-Rs (Fig. 1A). We found that 51% of pyramidal neurons (28/55) increased their firing during SPW-Rs (“SPW-R-active neurons”), while 11% of pyramidal cells (6/55) showed a decrease in firing (“SPW-R-suppressed neurons”) (Fig. 1B). For putative interneurons, 41% (28/68) and 18% (12/68) showed an increase and a decrease of their firing rates, respectively (Fig. 1B). In the lSuM, 24% of neurons (19/80) showed a significant increase of firing during SPW-Rs (“SPW-R-active neurons”) (Fig. 1B). The majority of ISuM neurons (76%) did not display a change in firing during SPW-Rs.

**Figure 1:**
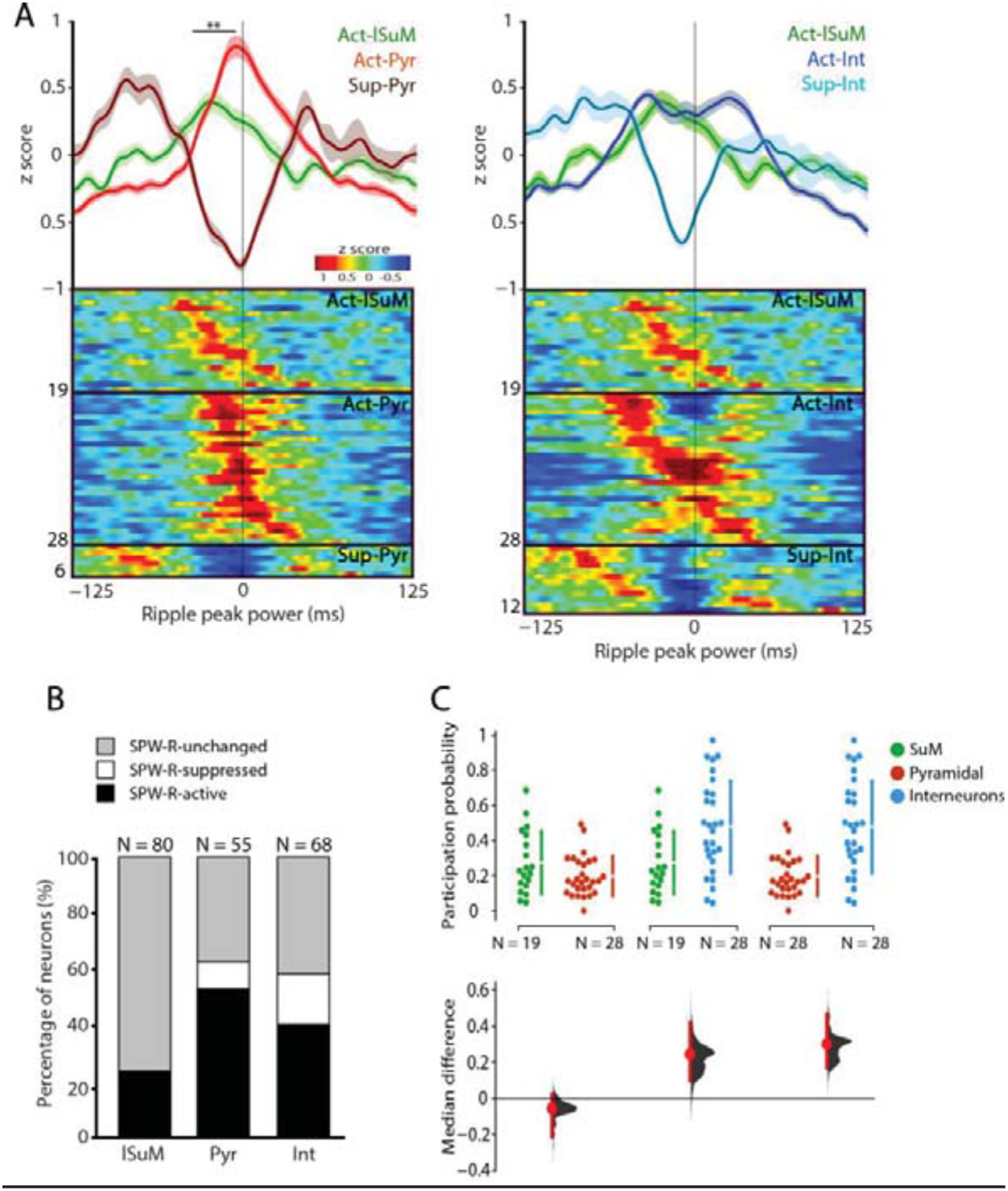
A population of lSuM neurons fire before hippocampal SPW-Rs. (A) Average (shaded error bars) firing curves per region and Z scored firing rate raster plots centered on SPW-R peak power for SPW-R-active and SPW-R-suppressed neurons. Each row represents the activity of a single neuron. Black vertical line indicates zero-time lag. Numbers to the left of raster plots represent the total number of neurons for every group. Left: Act-lSuM (lSuM SPW-R-active), Act-Pyr (putative hippocampal pyramidal SPW-R-active) and Sup-Pyr (putative hippocampal pyramidal SPW-R-suppressed). Right: Act-lSuM (lSuM SPW-R-active), Act-Int (putative hippocampal interneurons SPW-R-active) and Sup-Int (putative hippocampal interneurons SPW-R-suppressed). (B) Percentage of lSuM, pyramidal (Pyr) and interneurons (Int) classified as SPW-R-active, SPW-R-suppressed and SPW-R-unchanged. (C) Participation probability for lSuM neurons (n=19), pyramidal cells (n=28) and interneurons (n=28) during SPW-Rs, defined as the number of SPW-Rs in which a neuron fires divided by the total number of SPW-Rs. Interneurons show the highest participation probability as compared to lSuM and pyramidal neurons. lSuM shows a higher participation probability as compared to pyramidal neurons. The upper plots show the participation probability for each neuron. Vertical lines to the right of each group represent mean (gap in the lines) ± standard deviation (gapped lines). Lower plot indicates the median differences between groups. Each median difference in changes in participation probability is plotted as a bootstrap sampling distribution. Median differences are depicted as dots; 95% confidence intervals are indicated by the ends of the vertical error bars. Note the large dispersion of participation probabilities, from 0 to 0.7 for all cell types, and the presence of an important population of CA1 interneurons with high (>0.7) participation probabilities.

We then looked at the timing of the peak firing rate with respect to the SPW-R peak power (Fig. 1A). The peak firing rate occurred earlier than the SPW-R peak in most SPW-R-active neurons. The earliest firing neurons were the lSuM SPW-R-active neurons. The median time of peak firing was 19.2 ms before the SPW-R peak. The median time of firing suppression peak was 11 ms for SPW-R-suppressed interneurons and 4 ms for SPW-R-suppressed pyramidal neurons. The median time of peak firing was 6 ms for SPW-R-active interneurons and 2 ms for SPW-R-active pyramidal neurons. Significant differences were found for the following comparisons: lSuM and SPW-R-active pyramidal neurons (median difference= 17.7 ms [95.0% CI 3.6, 29.8], p= 0.00449, Kruskal test), SPW-R-active pyramidal and SPW-R-suppressed interneurons (median difference= −9.5 ms [95.0% CI −17.0, −3.0], p=0.00378, Kruskal test), and SPW-R-suppressed pyramidal and SPW-R-suppressed interneurons (median difference= −7.0 ms [95.0% CI −7.0, −7.0], p=0.0385, Kruskal test).

We then calculated the SPW-R participation probability for each neuron, which was defined as the number of SPW-Rs in which a neuron fired divided by the total number of SPW-Rs (Fig. 1C). Interneurons fired more frequently than lSuM neurons (median probability 0.47 and 0.23 for interneurons and lSuM neurons, respectively, median difference=0.24, [95.0% CI 0.0969, 0.422], p=0.00792, Kruskal test). Interneurons also fired more frequently than pyramidal neurons (median=0.17, median difference=0.301 [95.0% CI 0.167, 0.469], p=2.19e-05, Kruskal test). The participation probability of lSuM neurons tended to be higher as compared to pyramidal neurons, although the difference was not statistically significant (0.23 and 0.17 respectively, median difference= −0.0569 [95.0% CI −0.216, 0.023], p=0.165, Kruskal test). The main difference between CA1 interneurons and CA1 pyramidal/ISuM cells stems from the presence of highly participating interneurons showing a high probability (>0.7) to be active during SPW-Rs. Interestingly, the 3 groups have a widespread dispersion of values between 0 (cells that did not participate in SPW-Rs) and 0.7 (Fig. 1C). Replication studies are necessary to determine whether the cells can be divided in high and low participation probabilities.

### Firing properties of lSuM SPW-R-active neurons

We then assessed whether lSuM SPW-R-active neurons had different firing properties from SPW-R-unchanged neurons. To this aim, we compared several parameters characterizing the firing properties of the neurons during THE and SO states.

We found that SPW-R-active neurons had a higher firing rate during THE and SO than SPW-R-unchanged neurons: median=7.77 and 2.72 Hz for THE, respectively, median difference= −5.05 Hz [95.0% CI −8.53, −0.525], p=0.017, Kruskal test; and median=4.17 and 1.99 Hz for SO, respectively, median difference= −2.17 Hz [95.0% CI −8.88, −0.624], p=0.0119, Kruskal test, Fig. 2A left). Although some SPW-R-active and SPW-R-unchanged neurons increased or decreased their firing rate between THE and SO, at the population level the median firing rates were not significantly different between THE and SO: median difference= −1.79 Hz for SPW-R-active neurons [95.0% CI −7.04, 0.856], p=0.107, Wilcoxon test; median difference= −0.306 Hz for SPW-R-unchanged neurons [95.0% CI −1.49, 0.827], p=0.0247, Wilcoxon test (Fig. 2A right). Using the firing rate and spike properties (asymmetry and width), we did not find an obvious way to distinguish SPW-R-active from SPW-R-unchanged neurons (Figure 7C).

**Figure 2:**
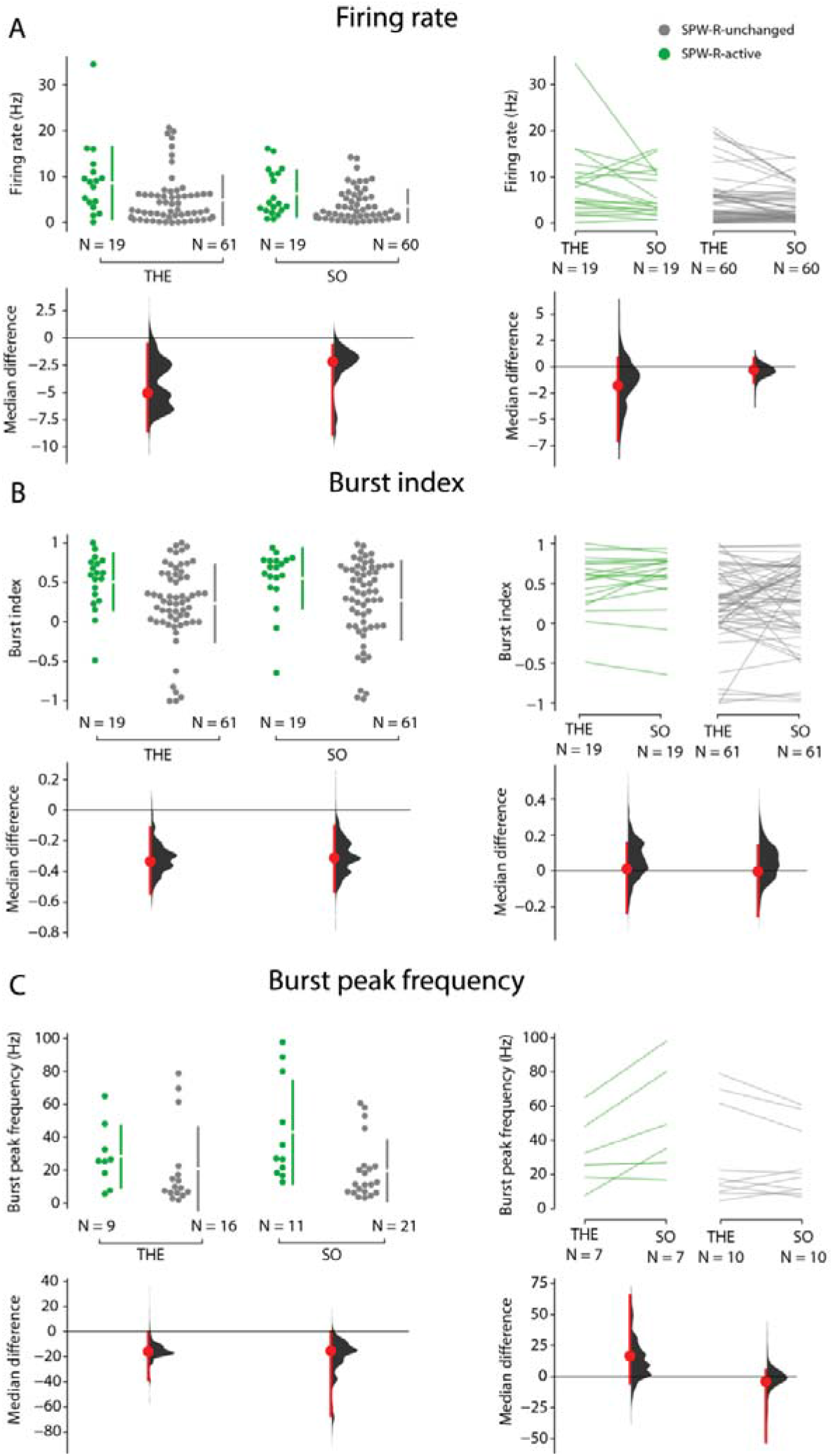
Firing rate and burst parameters for lSuM neurons during theta (THE) and slow oscillations (SO). (Left) In panels A, B and C, upper plot shows values per neuron, represented by dots. Vertical lines to the right of each group represent mean (gap in the lines) ± standard deviation (gapped lines). (Right) In panels A, B and C, upper plot shows values per neuron during THE and SO, represented by lines. Lower plot indicates the median differences between groups. Each median difference is plotted as a bootstrap sampling distribution. Median differences are depicted as dots; 95% confidence intervals are indicated by the ends of the vertical error bars. Firing rate (A left), burst index (B left) and burst peak frequency (C left) differences between SPW-R-active neurons (green) and SPW-R-unchanged neurons (grey). Firing rate and burst index was higher for SPW-R-active neurons during THE and SO. Burst peak frequency was higher for SPW-R active neurons only during SO. Firing rate (A right), burst index (B right) and burst peak frequency (C right) differences between THE and SO for SPW-R-active neurons (green) and SPW-R-unchanged neurons (grey). We did not find differences between THE and SO for either of the neuronal groups.

SPW-R-active neurons had a higher bursting activity than SPW-R-unchanged neurons during THE and SO. For THE: burst index median=0.59 and 0.26, respectively, median difference= −0.33 [95.0 % CI −0.547, −0.112], p=0.0151, Kruskal test; for SO: burst index median=0.69 and 0.38, respectively, median difference= −0.31 [95.0 %CI −0.532, −0.104], p=0.0144, Kruskal test, Fig. 2B left). Burst indices were similar between THE and SO: median difference=0.0109 for SPW-R-active neurons, [95.0% CI −0.232, 0.154], p=0.314, Wilcoxon test; median difference= −0.00227 for SPW-R-unchanged neurons [95.0%CI −0.25, 0.142], p=0.544, Wilcoxon test (Fig. 2B right).

The peak frequency during the bursting periods (intra-burst frequency) of the SPW-R-active neurons was similar to that of SPW-R-unchanged neurons during THE: median=25.59 and 9.54 Hz, median difference= −16.0 Hz [95.0% CI −38.7, −0.964], p=0.101, Kruskal test (Fig. 2C left). However, the intra-burst frequency was higher in SPW-R-active neurons than in SPW-R-unchanged neurons during SO: median=27.08 and 11.52 Hz, median difference= −15.6 Hz [95.0% CI −67.4, −0.174], p=0.00657, Kruskal test (Fig. 2C left). We found no significant differences between THE and SO for either of the neuronal groups. For SPW-R-active neurons: median=25.6 and 35.35 Hz for THE and SO, respectively, median difference=16.5, [95.0% CI −5.58, 65.3], p=0.0425, Wilcoxon test; for SPW-R-unchanged neurons: median=16.02 and 19.53 Hz for THE and SO, respectively, median difference= −3.94, [95.0% CI −52.6, 5.11], p=0.285, Wilcoxon test, Fig. 2C right). Interestingly, the display of all data points seems to indicate the existence of two clusters: one with low and one with high burst peak frequency in all conditions.

Autocorrelations analysis showed that the discharge probability peak occurred earlier in SPW-R-active than in SPW-R-unchanged neurons (Fig. 3A left). For THE: median=4 and 55 seconds, median difference=51.0 [95.0% CI 13.0, 1.12e+02], p=0.00145, Kruskal test; for SO: median=3 and 22.5 seconds, median difference=19.5 [95.0% CI 6.5, 49.0], p=5.73e-05, Kruskal test. This finding indicates that SPW-R-active neurons fire with short inter-spikes intervals, in line with the higher burst activity found in this group, as compared to SPW-R-unchanged neurons. Although some SPW-R-active and SPW-R-unchanged neurons strongly increased or decreased their time of discharge probability peak between THE and SO, at the population level there were no significant differences (Fig. 3A right). For SPW-R-active neurons: median difference=0.0 [95.0% CI −5, 1.0], p=0.375, Wilcoxon test; for SPW-R-unchanged neurons: median difference= −3.0 [95.0% CI −12.0, 23.0], p=0.049, Wilcoxon test.

**Figure 3:**
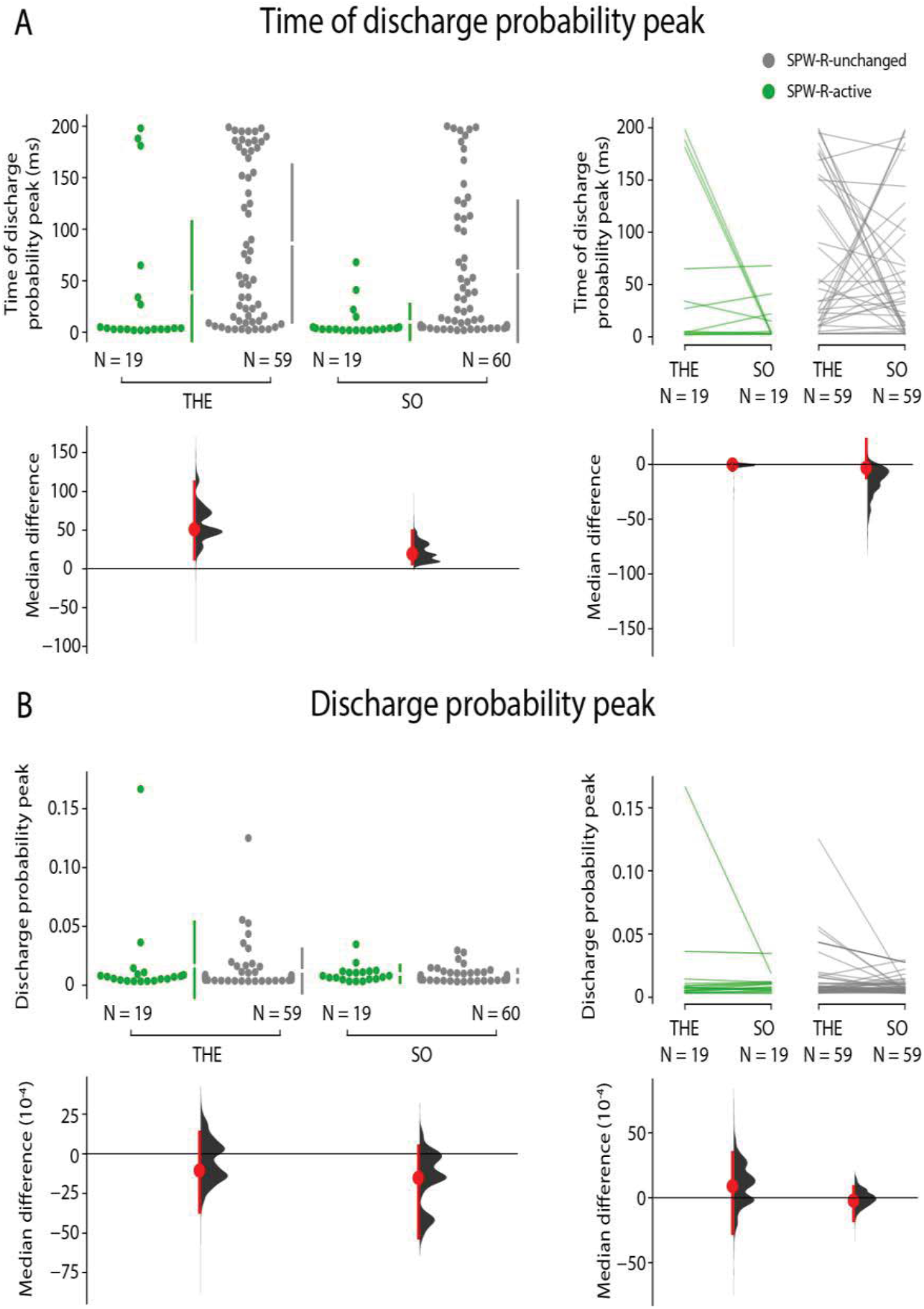
Firing parameters for lSuM neurons during theta (THE) and slow oscillations(SO). Time of discharge probability peak (A left) and discharge probability peak (B left) differences between SPW-R-active neurons (green) and SPW-R-unchanged neurons (grey). Discharge probability peak occurred earlier in SPW-R-active neurons during THE and SO. We did not find differences for discharge probability peak. Time of discharge probability peak (A right) and discharge probability peak (B right) differences between THE and SO for SPW-R-active neurons (green) and SPW-R-unchanged neurons (grey). We did not find differences between THE and SO for either of the neuronal groups. Same representation as in Figure 2.

The discharge probability peak was not statistically different between SPW-R-active and SPW-R-unchanged neurons (Fig. 3B left). For THE: median=0.0069 and 0.0059, respectively, median difference= −0.0011 [95.0% CI −0.0037, 0.0014], p=0.82, Kruskal test; for SO: median difference= −0.0015 [95.0% CI −0.0053, 0.00050], p=0.297, Kruskal test. We did not find differences in discharge probability peak between THE and SO for either of the two groups (Fig. 3B right). For SPW-R-active neurons: median difference=0.00089 [95% CI −0.0028, 0.0035], p=0.355, Wilcoxon test, for SPW-R-unchanged neurons: median difference= −0. 00022 [95%CI −0.0018, 0.00086], p=0.473, Wilcoxon test.

Altogether, these results show that lSuM SPW-R-active neurons have a greater firing activity as compared to SPW-R-unchanged neurons. This could be due to a global higher firing rate. We thus tested whether SPW-R-active neurons were also more active outside SPW-R periods. To this aim, we computed the firing rates of both types of neurons during ripples (SPW-R firing rate) and during SO periods excluding ripples times (i.e. inter-SPW-R firing rate). SPW-R-active neurons had a higher firing rate during both SPW-R and inter-SPW-R periods than SPW-R-unchanged neurons (Fig. 4A left). For SPW-R periods: median=6.5 and 1.93 Hz, median difference= −4.57 Hz [95.0 % CI −15.1, −2.74], p=5.05e-05 Kruskal test; for inter-SPW-R periods : median=3.62 and 1.6 Hz; median difference= −2.02 Hz [95.0% CI −9.21, −0.831], p=0.00259, Kruskal test. For SPW-R-active neurons, the firing rate was higher during SPW-R than during inter-SPW-R periods: median difference= −3.52 Hz [95.0% CI −12.3, −2.06], p=0.00549, Wilcoxon test (Fig. 4A right). However, there were no significant differences between SPW-R and inter-SPW-R epochs for SPW-R-unchanged neurons: median difference= −0.285 Hz [95.0% CI −1.53, 0.253], p=0.00219, Wilcoxon test (Fig. 4A right).

**Figure 4:**
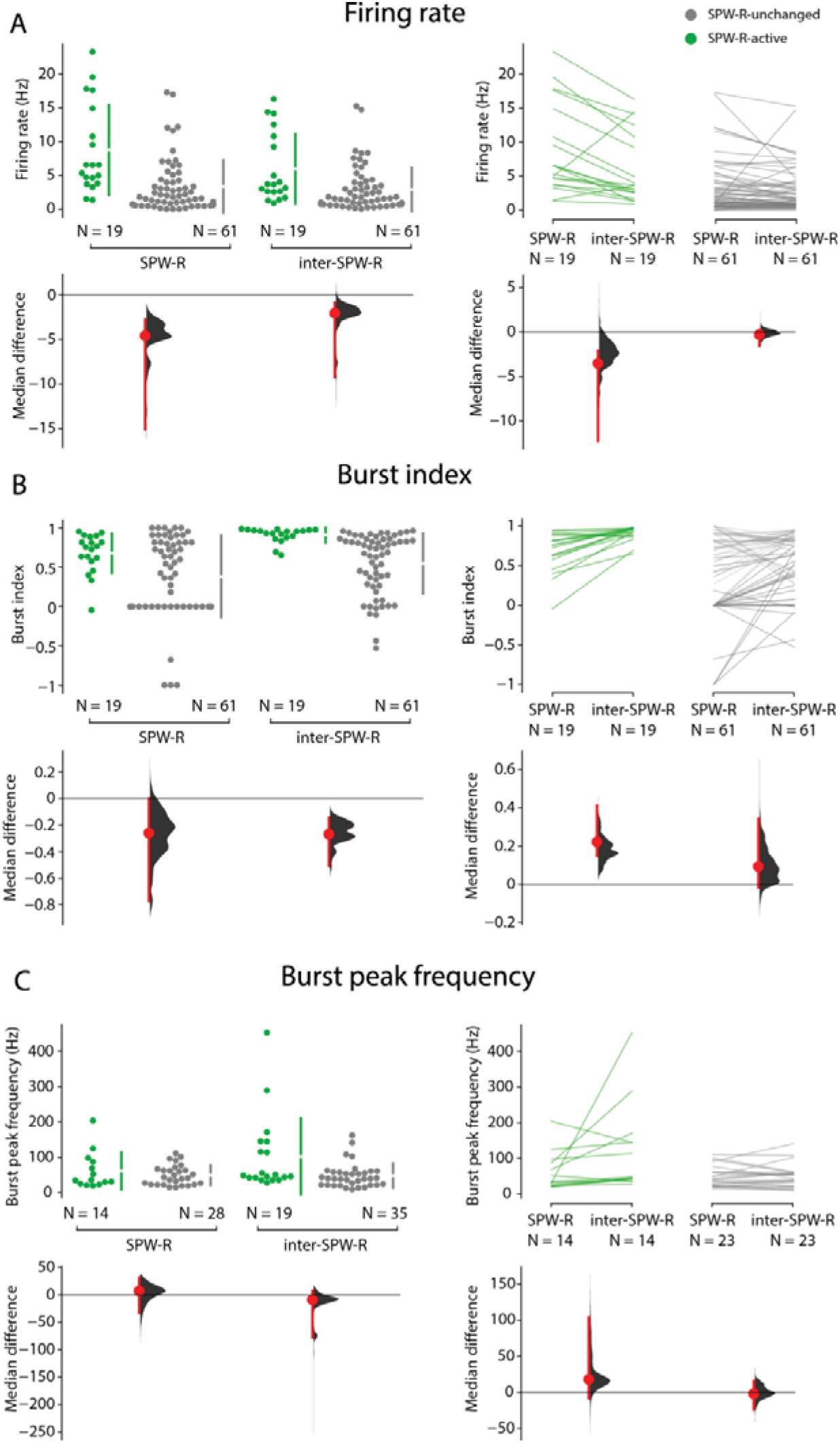
Firing rate and burst parameters for lSuM neurons during SPW-R and inter-SPW-R periods. Firing rate (A left), burst index (B left) and burst peak frequency (C left) differences between SPW-R-active neurons (green) and SPW-R-unchanged neurons (grey) during SPW-R and inter-SPW-R periods. Firing rate and burst index was higher for SPW-R-active neurons during SPW-R and inter-SPW-R periods. We did not find differences regarding burst peak frequency for either of the neuronal groups. Firing rate (A right), burst index (B right) and burst peak frequency (C right) differences between SPW-R and inter-SPW-R periods for SPW-R-active neurons (green) and SPW-R-unchanged neurons (grey). Firing rate was higher during SPW-R periods only for SPW-R-active neurons. Burst index was higher during inter-SPW-R periods only for SPW-R-active neurons. We did not find differences regarding burst peak frequency for either of the neuronal groups. Same representation as in Figure 2.

A higher firing rate for SPW-R-active neurons during SPW-Rs as compared to outside SPW-Rs, or as compared to SPW-R-unchanged neurons, may be due to increased bursting activity. We thus computed the burst index of both neuron populations for SPW-R and inter-SPW-R periods. Burst index was significant higher for SPW-R-active neurons only during inter-SPW-R periods (Fig. 4B left). For SPW-R periods: median=0.76 and 0.5 for SPW-R-active and SPW-R-unchanged neurons, respectively, median difference= −0.258 [95.0% CI −0.768, 0.0], p=0.0672, Kruskal test. For inter-SPW-R periods: median=0.96 and 0.69 for SPW-R-active and SPW-R-unchanged neurons, respectively, median difference= −0.266 [95.0% CI −0.506, −0.141], p=8.99e-07, Kruskal test (Fig. 4B left). When comparing burst indices between SPW-R and inter-SPW-R periods, we found a higher burst index during inter-SPW-R periods only for the SPW-R-active population (Fig. 4B right). For SPW-R-active neurons: median difference=0.22 [95.0% CI 0.149, 0.414], p=0.000293, Wilcoxon test; for SPW-R-unchanged neurons: median=0.0093 [95%CI −0.018 0.34], p=0.00423, Wilcoxon test.

Regarding intra-burst frequency, we found no significant differences between SPW-R-active and SPW-R-unchanged neurons either for SPW-R or for inter-SPW-R periods (Fig. 4C left). For SPW-R periods: median=35.09 and 42.29 Hz, 7.19 [95.0% CI −33.0, 31.1], p=0.689, Kruskal test; for inter-SWP-R periods: median=47.92 and 39.26 Hz for inter-SPW-R periods, −8.66 [95.0% CI −77.6, 7.28], p=0.0219, Kruskal test. We did not find significant differences when comparing SPW-R and inter-SPW-R periods (Fig. 4C right). For SPW-active neurons: mean=35.09 and 46.61 Hz for SPW-R and inter-SPW-R periods, respectively, median difference=18 [95.0 % CI −7.97, 104], p=0.0303, Wilcoxon test; for SPW-R-unchanged neurons: median=41.09 and 43.39 Hz for SPW-R and inter-SPW-R periods, respectively, median difference= −1.99 [95.0% CI −23.5, 15.4], p=0.976, Wilcoxon test.

We then examined how neuronal firing was modulated by the ongoing brain oscillation. Most of lSuM neurons were entrained by the ongoing oscillatory activity. Specifically, 93% (74/80) were entrained during THE epochs; and 98% (78/80) during SO epochs, p<0.05, Rayleigh test, Figure 7A). Whereas SPW-R-active and SPW-R-unchanged neurons did not display any difference in their theta-phase preference (median=191.64 and 168.9 °, respectively, median difference= −22.6 °, [95.0 % CI −57.8 °, 19.1 °], p=0.58, Kruskal test), SPW-R-active population fired later during the UP state of the slow oscillation as compared to SPW-R-unchanged population during SO (median=220.42 and 203.59 °, respectively, median difference= −16.9 °, [95.0 % CI −31.0 °, −8.87 °], p=0.0093, Kruskal test)(Figure 7A). Although we found significant differences between the two neuronal populations during SO, similar distribution of the individual values point to a weak significant difference (Figure 7B). We did not find differences for the strength of the entrainment between the two populations either for THE or for SO (median=0.17 and 0.15 for THE, median difference= −0.016 [95.0% CI −0.0786, 0.0575], p=0.91, Kruskal test; median=0.21 and 0.19 for SO, median difference= −0.0311 [95.0% CI −0.0863, 0.0555], p=0.88, Kruskal test).

Combining all previous results, we show that it is possible to clearly separate SPW-R-active from SPW-R-unchanged neurons on the basis of their firing properties (Fig. 5).

**Figure 5:**
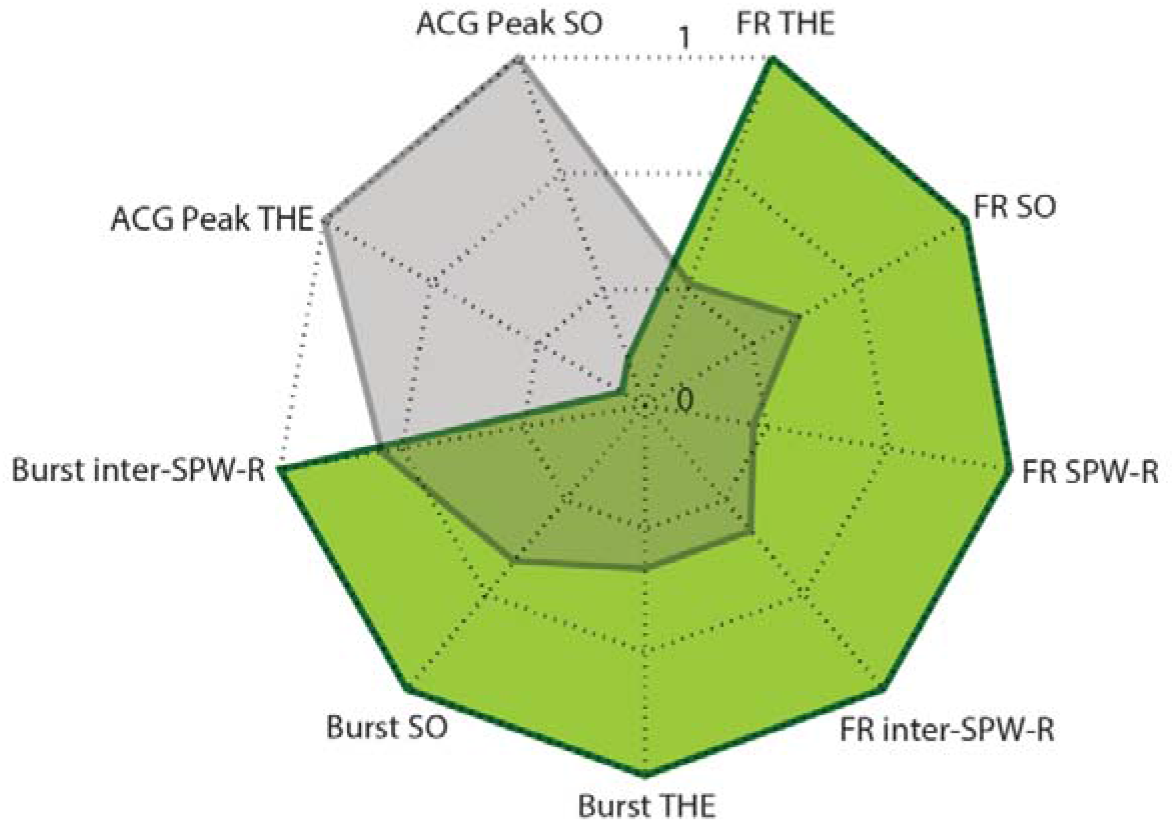
Radar plot of firing parameters showing significant differences between lSuM SPW-R-active (green) and SPW-R-unchanged neurons (grey). Every axis represents a firing parameter and its corresponding mean value for SPW-R-active and SPW-R-unchanged neurons. All values corresponding to the same group are connected to form a polygon. All values are normalized. The relative position and angle of the axes are not informative. FR THE and FR SO: firing rate during theta (THE) and slow oscillation (SO), FR SPW-R and inter-SPW-R: firing rate during SPW-R and inter-SPW-R periods, Burst THE and SO: burst index during THE and SO, Burst inter-SPW-R: burst index during inter-SPW-R periods, ACG Peak THE and SO: time of discharge probability peak during THE and SO. The distinct shape of the polygons illustrates the differences between SPW-R-active and SPW-R-unchanged neurons according to their firing properties.

### Correlation between lSuM firing and SPW-R parameters

Finally, we examined whether the firing of both lSuM SPW-R-active neurons and SPW-R-unchanged neurons was correlated with SPW-R properties like their duration, peak power and frequency. We found a negative but weak correlation between SPW-R-active neurons firing and SPW-R duration (r= −0.12, [95.0% CI −0.17, −0.07], p=7.7597e-07, Pearson correlation, Fig. 6A left), whereas the correlation was positive for SPW-R-unchanged neurons (r=0.23, [95.0% CI −0.18, 0.28], p=5.3796e-21, Pearson correlation, Fig. 6A right). Regarding SPW-R peak power, SPW-R-active neurons firing showed a positive correlation (r=0.41, [95.0% CI 0.37, 0.41], p=7.5342e-68, Pearson correlation, Fig. 6B left) and SPW-R-unchanged neurons showed no correlation (r= −0.04, [95.0% CI −0.08, 0.01], p=0.15, Pearson correlation, Fig. 6B right). SPW-R-active neurons also displayed a negative correlation with SPW-R frequency (r= −0.07, [95.0% CI −0.12, −0.03], p=0.002, Pearson correlation, Fig. 6C left) while no correlation was found for SPW-R-unchanged neurons r=0.03, [95.0% CI −0.02, −0.08], p=0.22, Pearson correlation, Fig. 6C right). These results indicate that lSuM firing during SPW-Rs depends on some SPW-R parameters. SPW-R-active neurons are more active during high peak power SPW-Rs, whereas SPW-R-unchanged neurons are more active during long SPW-R.

**Figure 6:**
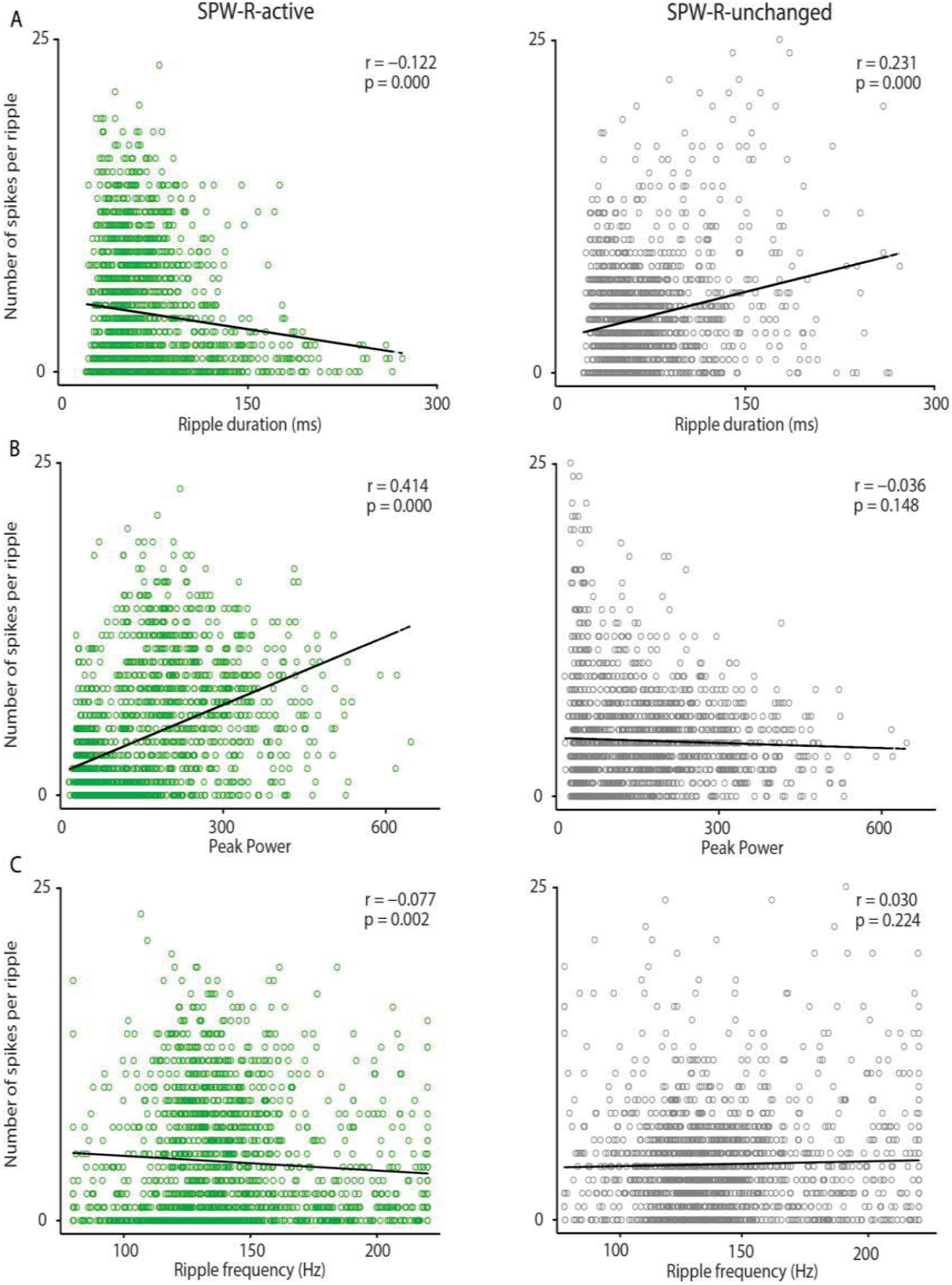
Correlation between ripples parameters and lSuM firing during ripples. Each circle indicates number of spikes per ripple for lSuM SPW-R-active (green, left) and SPW-R-unchanged neurons (grey, right), and its corresponding ripple parameter. Correlation coefficient (Pearson correlation) and p value are shown. (A) Ripple duration and number of spikes per ripples are negatively correlated for SPW-R active neurons and positively correlated for SPW-R unchanged neurons. (B) Ripple peak power and number of spikes are positively correlated for SPW-R active neurons. (C) Ripple frequency and number of spikes are negatively correlated for SPW-R active neurons.

**Figure 7:**
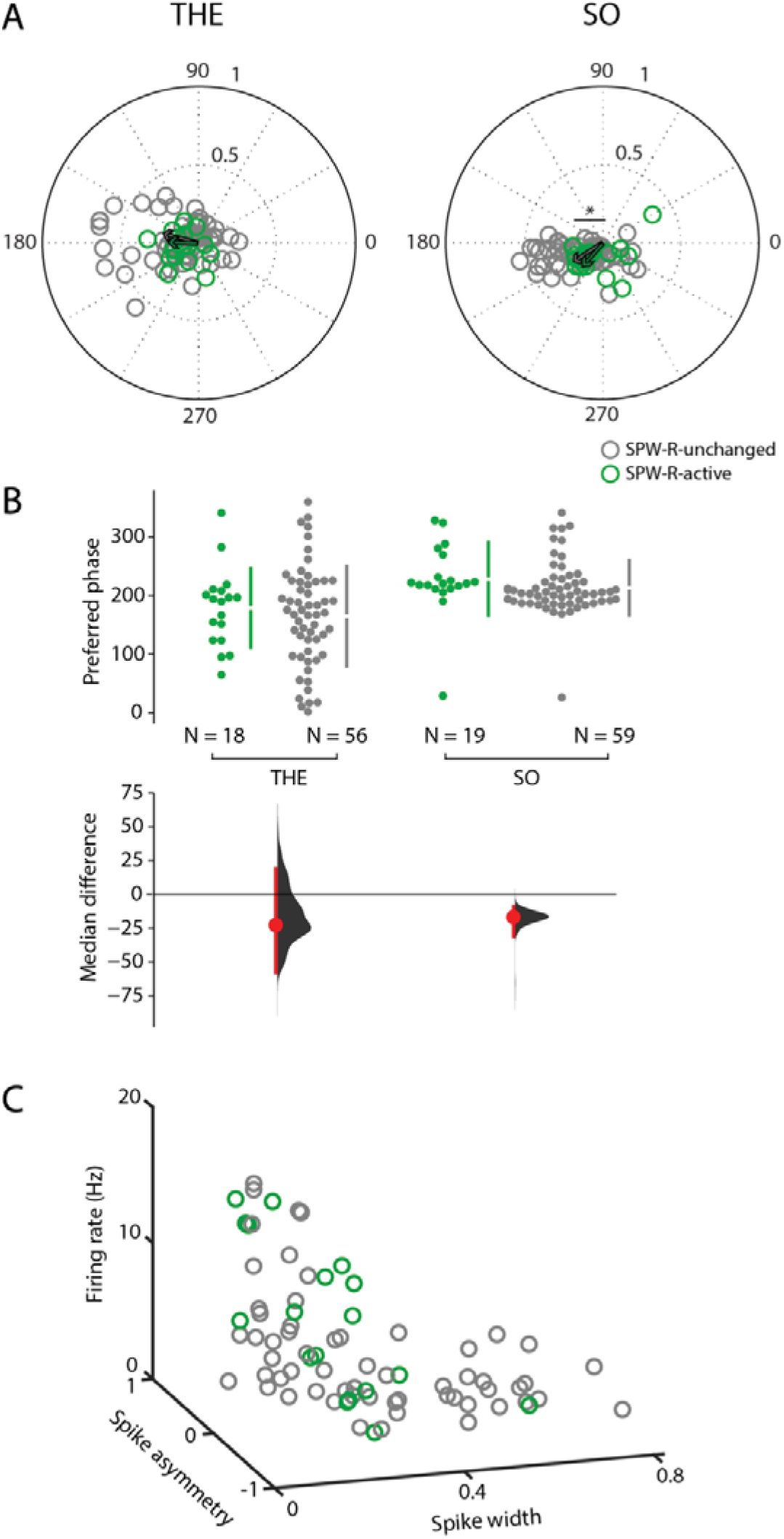
Entrainment of lSuM neurons by theta (THE) and slow oscillations (SO). (A) Polar plots of preferred phase and modulation strength of SPW-R active (green) and SPW-R-unchanged neurons significantly entrained (Rayleigh test, p<0.05) during THE (left) and SO (right). The green and grey arrows indicate the mean preferred phase and strength of modulation for the significantly entrained SPW-R-active and SPW-R-unchanged neurons, respectively. (B) Preferred phase differences between SPW-R-active (green) and SPW-R-unchanged neurons (grey) during THE (left) and SO (right). We found differences only during SO. Same representation as in Figure 2 left. (C) Relationship among trough-to-peak latency (spike width), asymmetry index of the spike waveform (spike asymmetry),and firing rate for Supramammillary nucleus and hippocampus interaction during ripples SPW-R-active (green) and SPW-R-unchanged neurons (grey). We could not distinguish between SPW-R-active from SPW-R-unchanged neurons.

## Discussion

The interaction between SuM and HPC has classically been focused in the context of hippocampal theta oscillations due to SuM connectivity with other nuclei involved in theta generation, such as the reticular formation and the septum (Green & Arduini, 1954). In the present study we characterized neuronal activity in the lSuM during both theta and slow oscillation-dominated states and its interaction with hippocampal SWP-Rs in anesthetized rats. We found SPW-R-associated neuronal firing in a population of lSuM neurons. The latter increased their firing prior SPW-R peak power and prior CA1 principal cells. They showed distinct firing properties and tend to fire more during high peak power ripples. These findings suggest the existence of an interaction between lSuM and hippocampal SPW-Rs and raise questions about the causality of this interaction and its functional significance.

We demonstrate a population of lSuM neurons that increase their firing around 20 ms before dorsal CA1 SPW-Rs, which suggests coordinated activity between both regions. However, the lSuM is not directly connected to dorsal CA1 HPC (Magloczky *et al.*, 1994; Vertes, 1992), where SPW-Rs were recorded. This lSuM-SPW-R interaction may be explained by indirect connections between lSuM and dorsal CA1. The lSuM innervates stratum oriens and pyramidal layer of dorsal CA2 and CA3 (Magloczky *et al.*, 1994; Vertes, 1992), hippocampal regions postulated to generate SPW-Rs (Oliva *et al.*, 2016; Csicsvari *et al.*, 2000). The lSuM also sends afferents to the dentate gyrus, which, in turn, connects to CA2 (Vertes & McKenna, 2000). Sullivan *et al.* (2011) proposed that dentate gyrus inputs modulate the occurrence of SPW-Rs as there is a positive correlation between dentate gamma power and the frequency of SPW-R occurrence. The lSuM has an excitatory effect on the dorsal dentate gyrus (Carre & Harley, 1991; Nakanishi *et al.*, 2001) but also an inhibitory effect (Segal, 1979; Mizumori *et al.*, 1989), likely due to the dual glutamatergic and GABAergic phenotype of lSuM projections to the dentate gyrus (Soussi *et al.*, 2010). Both excitatory and inhibitory lSuM projections to the dentate gyrus could be responsible of an indirect influence on SPW-Rs. Moreover, CA2, CA3 and the dentate gyrus connect dorsal CA1 (Amaral & Witter, 1989), which may also explain the origin of the CA1 SPW-R-associated neuronal firing found in the lSuM.

The lSuM-SPW-R related firing could also be explained by their reciprocal connections with the septum complex. The lSuM sends afferents to the medial and lateral septum contacting both principal cells and interneurons; and at least 25% of these projections connect to the dorsal HPC as well (Borhegyi *et al.*, 1998; Kiss *et al.*, 2000; Leranth & Kiss, 1996). Furthermore, Unal *et al.*, 2018 described two neuronal populations in the medial septum showing SPW-R related activity: one population suppressed its firing during SPW-Rs while the other population was activated. Finally, the septum sends projections to hippocampal interneurons (Freund & Antal, 1988). These results raise the possibility of an indirect pathway from the SuM to CA1 via the septum, which may explain the lSuM SPW-R related firing identified in the present work.

The firing of lSuM activated neurons precedes hippocampal neuronal firing during SPW-Rs. It would be tempting to propose that the lSuM may act as a hippocampal driver. However, caution is in order. If SPW-Rs are activities emerging from a large network of brain regions, early firing may just mark the onset of the emergence of the event, without necessarily implying causality. Nevertheless, lSuM firing is followed by SPW-R-active interneurons, and by SPW-R-active pyramidal neurons. Oliva *et al.* (2016) have also shown that CA1 interneurons fire before CA1 pyramidal neurons during SPW-Rs. In our experiments, lSuM neurons fire around 20 ms before SPW-R occurrence, approximately at the same time CA2 pyramidal neurons and interneurons recorded by Oliva *et al.* (2016) are firing. Thus, lSuM and CA2 neurons may simultaneously modulate CA1 activity. Circuit manipulation experiments using optogenetic tools are now needed to investigate in further details the role of lSuM in SPW-R and hippocampal firing modulation.

lSuM SPW-R-active neurons show different firing properties as compared to lSuM SPW-R-unchanged neurons. SPW-R-active neurons discharge more frequently and tend to burst during both THE and SO states. The intra-burst frequency is also higher for SPW-R-active neurons. The discharge probability peak from the autocorrelogram occurs earlier during both THE and SO for SPW-R-active neurons, which is in accordance with the higher burst activity of this population. SPW-R-active neurons are also more active than SPW-R-unchanged neurons during SPW-R periods, which is in accordance with their increase of firing during SPW-Rs. The fact that we also found differences between these neuronal groups during inter-SPW-R periods demonstrates the existence of two functionally distinct neuronal populations in lSuM: SPW-R-active neurons characterized by a higher excitability during THE and SO and an increase of firing during SPW-Rs, and SPW-R-unchanged neurons characterized by a lower excitability and no changes of firing during SPW-Rs. We hypothesize that SPW-R-active neurons correspond to SuM projections connecting both medial septum and dorsal HPC (Borhegyi *et al.*, 1998), which are likely to be glutamatergic/aspartatergic (Kiss *et al.*, 2000). These 3 areas, lSuM, medial septum and dorsal HPC, could be part of a circuit involved in SPW-R modulation.

SPW-Rs are considered an important mechanism to transfer memory from hippocampus to neocortex (Buzsáki, 2015). The fact that lSuM firing is coordinated with SPW-Rs suggests a role of this hypothalamic nucleus in memory. Supporting this hypothesis, Shahidi *et al.* (2004) found deficits in consolidation of a passive avoidance task when reversible inactivation of SuM was performed in rats. In addition, evidence of memory impairment after electrolytic lesion or inactivation of the SuM has been shown (Aranda *et al.*, 2006; Aranda *et al.*, 2008). Consistently with these findings, Ito *et al.* (2009) found an increase of c-fos expression in SuM after exploring a novel environment. In a recent work, Ito *et al.* (2018) showed that optogenetic inhibition of SuM alters its involvement in a working memory task. Although these works point to a role of SuM in memory, neither of them studied memory during sleep, period in which memory consolidation is supposed to occur (Rasch & Born, 2013). Studying the neuronal activity of SuM and HPC during the sleep period following the learning of a task would help us to understand if the lSuM-hippocampal interaction during SPW-Rs we described here reflects memory processes.

## Acknowledgements

The authors wish to thank support from FRM, FFRE, and CURE Epilepsy Taking Flight Award.

